# Small molecule modulates *α*-Synuclein conformation and its oligomerization via Entropy Expansion

**DOI:** 10.1101/2022.10.20.513005

**Authors:** Sneha Menon, Jagannath Mondal

**Affiliations:** Tata Institute of Fundamental Research, Center for Interdisciplinary sciences, Hyderabad 500046, India

## Abstract

Aberrant misfolding and progressive aggregation of the intrinsically disordered protein (IDP), ***α***-synuclein, are associated with the etiology of several neurodegenerative diseases. However, the structurally heterogeneous ensemble of this IDP and lack of a well-defined binding pocket make it difficult to probe the druggability of ***α***-synuclein. Here, by building a comprehensive statistical model of the fuzzy ensemble of a millisecond-long atomistic simulation trajectory of monomeric ***α***-synuclein interacting with the small-molecule drug fasudil, we identify exhaustive sets of metastable binding-competent states of *α*-synuclein. The model reveals that the interaction with the drug primes this IDP to explore both more compact and more extended conformational sub-ensemble than those in neat water, thereby broadening its structural repertoire in presence of small-molecule via an *entropy expansion* mechanism. Subsequent simulation of the dimerisation process shows that similar motif of entropic-expansion mechanism helps fasudil to retard the self-aggregation propensity of *α*-synuclein via trapping it into multiple distinct states of diverse compaction featuring aggregation-resistant long-range interactions. Furthermore, small-molecule binding interactions in dimerisation-competent relatively extended states have a screening effect that hinders the formation of stable dimer contacts. Together, the investigation demonstrates the ability of small-molecules to have an ensemble-modulatory effect on IDPs that can be effectively utilised in therapeutic strategies probing aggregation-related diseases.

## Introduction

Intrinsically disordered proteins (IDPs) comprise a class of proteins that lack a unique, well-defined three-dimensional structure under physiological conditions. Notwithstanding the structural disorder, these proteins can perform a variety of functions in a number of biological pathways thus challenging the classical structure-function paradigm. ^1,2^ Mainly due to their promiscuous binding nature, these IDPs are involved in crucial cellular processes such as regulation of cell cycle signaling and act as hubs in interaction networks.^3–5^ Emergent studies present disordered proteins as an important component of biomolecular condensates that play regulatory roles in several key cellular functions. ^6–9^

Considering the broad range of functionality of IDPs, their disregulation and altered abundance can lead to a number of diseases including cancer, diabetes, cardiovascular and neurodegenerative disorders. ^1,2,10^ In rational structure-based drug design, exploration of the target proteins is a crucial step based on which the interactions of small molecules are optimized to have stabilising interactions with a structured ligand-binding pocket.^11,12^ This traditional approach has mainly been developed considering folded proteins as the potential drug-target. IDPs, on the other hand, are considered *undruggable* since they exist as an ensemble of multiple distinct conformational states that are in dynamic equilibrium under physiological conditions. Therefore, due to lack of structured ligand-binding domains, IDPs are not amenable to traditional structure-based drug-design paradigm.^13^ Nevertheless, a few monomeric disordered proteins have been successfully targeted by small molecules. ^14^ Oncogenic transcription factor c-Myc has been explored for its small-molecule interaction.^15^ Abeta42 has remained a popular target system for small-molecule drugs. ^16–20^ In addition, other IDPs associated with neurodegenerative disorder such as Tau,^21^ KTBP,^22^ osteopontin^23^ have also been probed by small-molecule drug-discovery endeavours.

The IDP of this article’s interest, alpha-synuclein (*α*S) has remained an important drug-target^24–26^ owing to its long-rooted association with neurodegenerative diseases. Often, the early aggregation stages of *α*S into small oligomers is implicated in the causation of Parkinson’s disease^27,28^ and a potential therapeutic strategy has been the stabilization of *α*S in soluble monomeric form. In particular, the small molecule fasudil (focus of the present investigation) has been shown to interact with *α*S and retard its aggregation^26^ In this study, the anti-aggregative effect of fasudil on *α*S assembly was tested both in vivo and in vitro. Fasudil was found to significantly reduce *α*S aggregation in vitro and NMR spectroscopy revealed the direct interaction of fasudil with residues in the charged C-terminal of *α*S. Moreover, fasudil treatment significantly diminished *α*S pathology in vivo with improvement in cognitive and motor functions. Detailed characterisation of the mechanisms by which small molecules can interact with disordered proteins and potentially alter their disease promoting behaviour, could have profound implications in drug-discovery for disordered protein targets. However, the highly flexible and broad conformational ensemble of IDPs and the consequential challenges associated with conformational characterisation at atomic resolution makes it difficult to probe the druggability of IDPs using small molecules. Characterisation of the effect of small molecule binding on the ensemble properties of IDPs and the molecular details of the stabilizing interactions can provide the basis for selectivity and can be effectively calibrated to induce the desired effect. In particular, latest effort by Robustelli and coworkers^29^ had produced unprecedently long-time scale (1500 *µ*s) all-atom molecular dynamics (MD) simulation trajectory of small molecule drug fasudil with monomeric *α*S, in which the predicted protein-fasudil interactions were found to be in good agreement with previously reported NMR chemical shift data. However, a routine comparison of monomeric *α*S ensemble in neat water^30^ with that in presence of fasudil does not bring out significant difference, as customary signature of the dynamic nature of an IDP ensemble.

In the current work, we take an initiative of exhaustive dissection of the conformational ensemble of *α*S in water as well as in aqueous fasudil solution into key metastable sub-ensembles with an aim for atomistic characterisation of the action of small-molecule on *α*S. Towards this end, we investigate the large atomistically simulated ensemble of monomeric *α*S in neat water as well as in aqueous fasudil solution by developing individual Markov State Models (MSM) of this IDP in the two different media. Our investigation results show that, apart from partly retaining the prevalent sub-ensembles in water, the presence of fasudil gives rise to distinctly new sub-ensembles of monomeric *α*S via the *entropic expansion* mechanism.^20,31–33^ Characterisation of small-molecule interaction with the protein indicates that small-molecule fasudil has differential interactions with each of these macrostates that traps the protein conformations in these states thus expanding its conformational entropy. To explore how the ensemble modulatory effect of fasudil translates to its ultimate role in aggregation, we simulated the effect of fasudil on *α*S dimerisation process. The presence of fasudil in the solution indeed significantly hindered the early dimerisation of *α*S due to its preferential interactions with the protein residues mainly in the charged C-terminal. The dimerising ensemble in fasudil solution corroborates with the entropy expansion mechanism as seen in the monomer ensemble.

## Results and Discussion

### 1. Presence of small-molecule retains inherent fuzziness in the ensemble of *α*S monomer

Ultra long and atomistically modelled MD simulation trajectories of monomeric *α*S in water and in aqueous fasudil media formed the base of the current investigation. In particular, we utilised 108 *µ*s-long ensemble of *α*S in water, combinedly simulated in Anton supercomputers^30^ and in-house GPU-based servers. On the other hand, a 1500 *µ*s-long ensemble of monomeric *α*S in a solution of fasudil, generated in Anton supercomputer^29^ was employed in the current investigation. Visual inspection of the ensemble of *α*S in neat water vis-à-vis the ensemble in aqueous fasudil solution provided an impression of a highly heterogeneous mixture of a large number of conformations, a signature of intrinsic disorder inherent in *α*S (see Figure 1a). More significantly, in a clear departure from the classical view of ligand binding to a folded globular protein, the visual change in *α*S ensemble due to the presence of small molecule is not so strikingly apparent. As a result, we surveyed for a suitable quantitative metric to characterise the potential change in conformations of this IDP in presence of the small molecule fasudil. For identifying an appropriate collective variable (CV) that can be used to distinguish between the different metastable states of the system and their transitions in a more optimal fashion, we quantified the relative slowness of a set of candidate features that are commonly used for IDPs via a novel feature-scoring method called VAMP2 score.^34,35^ As introduced by Noé and coworkers, VAMP2 score is based on the Variational Approach for Markov Processes (VAMP) and maximizes the kinetic variance in the features.^34,35^ Therefore, the VAMP2 score can be used to compare the effectiveness of a set of features to capture the slow dynamics of the biomolecular system. The candidate CVs of our choice were namely, radius of gyration (R_*g*_), sets of all inter-residue contact probabilities and sets of all backbone torsions present in *α*S. Very interestingly, assessment of VAMP2-score for both the ensembles (in neat water as well as in presence of aqueous fasudil solution) across these CVs (Figure S1a,b) indicated that R_*g*_ remains the slowest varying metric for *α*S in both water and aqueous fasudil solution. In hindsight, this observation justifies the regular usage of R_*g*_ as a key metric in many experimental measurements pertaining to IDPs. Accordingly, we proceeded with R_*g*_ to compare the conformational space of *α*S in water versus aqueous fasudil solution. However, comparison of the probability distribution of R_*g*_ of the full ensembles (Figure 1b) showed only small differences between water versus aqueous fasudil solution. The distribution of the ensemble in water is slightly broader with the peak shifted towards higher value of R_*g*_ (peak position at 2.22 nm and 2.10 nm in neat water versus in aqueous fasudil solution), suggesting that this ensemble has a relatively higher population of extended conformations compared to the ensemble in aqueous fasusdil solution. However the small difference in the distribution indicates that a whole-ensemble view of the conformations does not sufficiently distinguish *α*S in neat water from that in aqueous fasudil solution. This also suggested that a more exhaustive state-space decomposition of the conformational ensemble of the IDP is required for gleaning into the role of the small-molecule in modulating the ensemble of *α*S.

**Figure 1:**
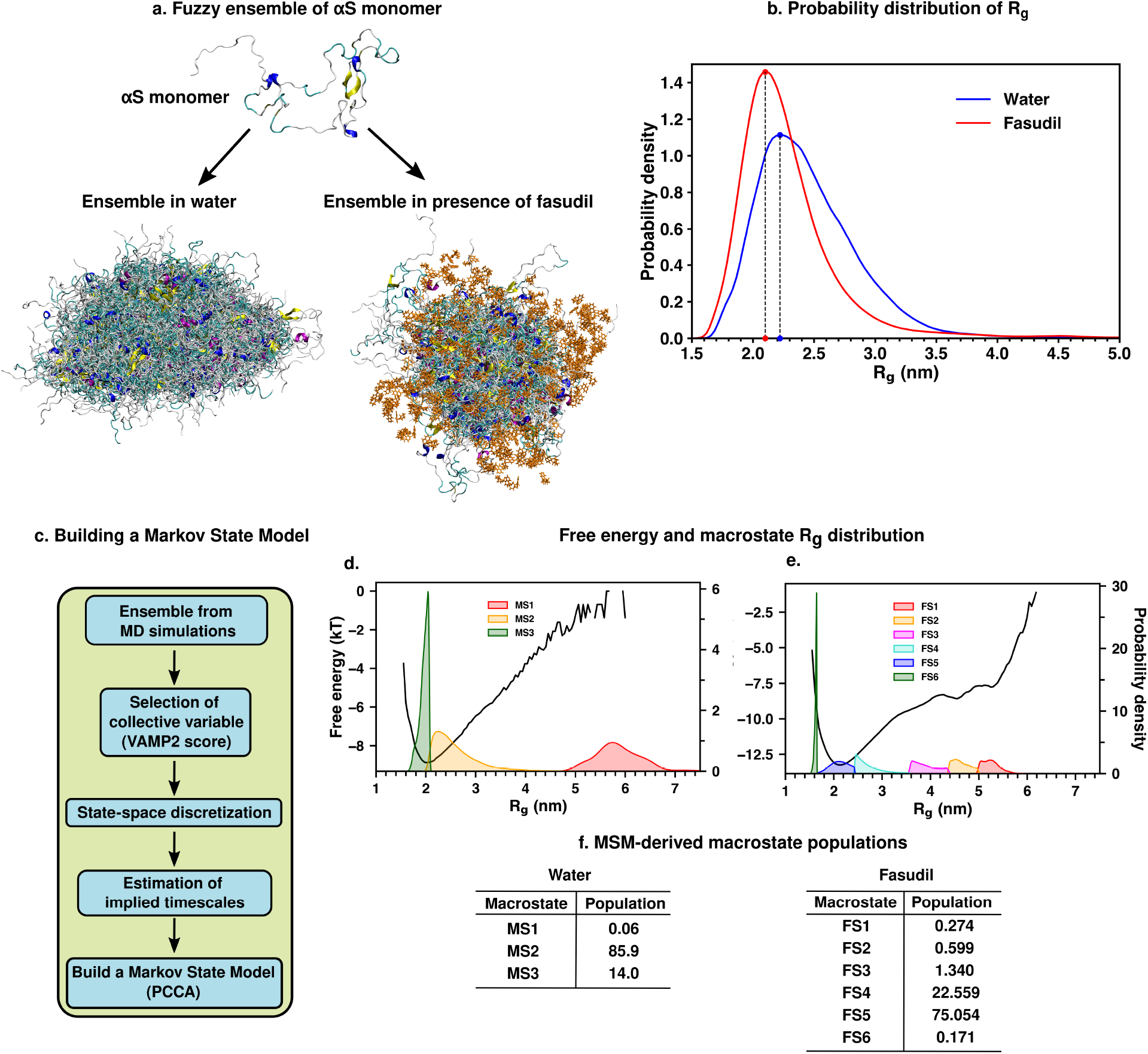
(a) A monomeric conformation of *α*S monomer is shown. A fuzzy ensemble of *α*S monomer in water and aqueous fasudil solution is presented. The protein is colored by secondary structure and the fasudil molecules are shown in orange (b) Probability distribution of R_*g*_ of the two *α*S ensembles. (c) A flowchart of the process of building a Markov State Model. Probability density of the macrostates obtained from the MSM corresponding to the ensemble in (d) water and (e) aqueous fasudil solution, overlaid with the one-dimensional free-energy landscape estimated as a function of R_*g*_. (f) The percentage macrostate populations in neat water and aqueous fasudil solution.

### 2. Markov State Model elucidates distinct binding competing states of *α*S in presence of drug molecules

For identifying key conformational sub-ensembles of *α*S in water and in aqueous fasudil solution, we undertook the development of a Markov State Model (MSM) by kinetically clustering the large atomistic simulation trajectories in these two respective environments. Towards this end, the slowest varying feature in both the ensembles (in presence or absence of fasudil) i.e. R_*g*_ of *α*S was used to discretise the time-continuous trajectory as a first step of the MSM. Analysis of the implied time scales as a function of lag-time (Figure S2) predicted a Markovian model of *α*S in water with three temporally separated states (see Figure 1f for relative population). On the other hand, a similar analysis of *α*S in presence of fasudil clearly indicated the distinct presence of spatially and temporally resolved six conformational metastable states (see Figure 1f for relative populations). The probability densities of these macrostates (depicted in Figure 1d, e) along R_*g*_ space indicate that these states are spatially non-overlapping and mutually distinct.

A pertinent question arises: Are the appearances of newer states in *α*S in presence of fasudil a result of the presence of longer trajectory in fasudil solution (1500 *µ*s) than that compared to the ensemble generated in neat water (108 *µ*s)? We verified this by building an MSM for blocks of data of fasudil system : using 108 *µ*s (same as in water system) and 700 *µ*s segment of the trajectory (see Figure S3). Very interestingly, the MSM built using lesser data (and same amount of data as in water) also indicated the presence of six states of *α*S in presence of fasudil, as was observed in the MSM of the full trajectory. Together this exercise invalidates the sampling argument and suggests that the increase in the number of metastable macrostates of *α*S in fasudil solution relative to that in water is a direct outcome of the interaction of *α*S with the small molecule.

A closer comparison of the mutual correspondence between the metastable states of *α*S in water versus those in the presence of fasudil along R_*g*_ space indicates that while fasudil dissects the states present in water further, it also creates more compact as well as more extended states than those favored in water (Figure 1d,e). Hereafter, we will refer to macrostates created in presence of fasudil prefixed with FS and those generated in water prefixed with MS. The most dominant macrostates in fasudil (FS4 and FS5) jointly correspond to the most populated sub-ensemble (MS2) of *α*S in water. Similarly, the most extended state in fasudil solution FS1 approximately corresponds to most extended state (MS1) in water. However, the comparison of mean R_*g*_ (FS1: 5.255 (*±* 0.179 nm) vs MS1: 5.859 (*±* 0.437 nm)) suggests that this state is more extended in water. Finally, comparison of R_*g*_ space of the most compact states in water (MS3) shows close correspondence to a subset of one of the most compact states *α*S in fasudil (FS5). One of the most intriguing observations of the current investigation is that apart from populating the aforementioned metastable states similar to that in water, the interaction of fasudil with *α*S introduces three completely new states namely, FS2, FS3 and FS6. In particular, FS2 and FS3 are extended states having R_*g*_ distribution intermediate between the dominant compact states (FS4 and FS5) and the most extended state FS1. On the other hand, FS6 is the most compact state resulting from fasudil-*α*S interaction. The comparison of average R_*g*_ indicates that this new state is more compact than MS3, the most compact state in water.

### 3. Fasudil exhibits conformation-dependent interactions with individual metastable states

The diverse extent of compaction across the key macrostates in presence of fasudil suggested that the interaction of *α*S with small-molecules in different conformational states might be a key factor in governing the conformational features. While precedent investigations^26,29^ have grossly reported dominant fasudil-*α*S interaction via the C-terminal region of the protein, we predicted that a more quantitative analysis characterising fasudil’s residue-wise interaction profiles with each of the conformational macrostates would provide atomistic insights on fasudil’s manner of modulation of individual conformational macrostates. Accordingly, we calculated the contact probabilities between fasudil and each *α*S residue. A contact between fasudil and a protein residue is considered to be present when the minimum distance between any heavy atom of fasudil and the protein is less than 0.6 nm. This is calculated for the entire population of each macrostate to determine the contact probabilities. The *α*S:fasudil contact maps for the six macrostates are presented in Figure 2a and snapshots of the interactions are presented in Figure 2b.

**Figure 2:**
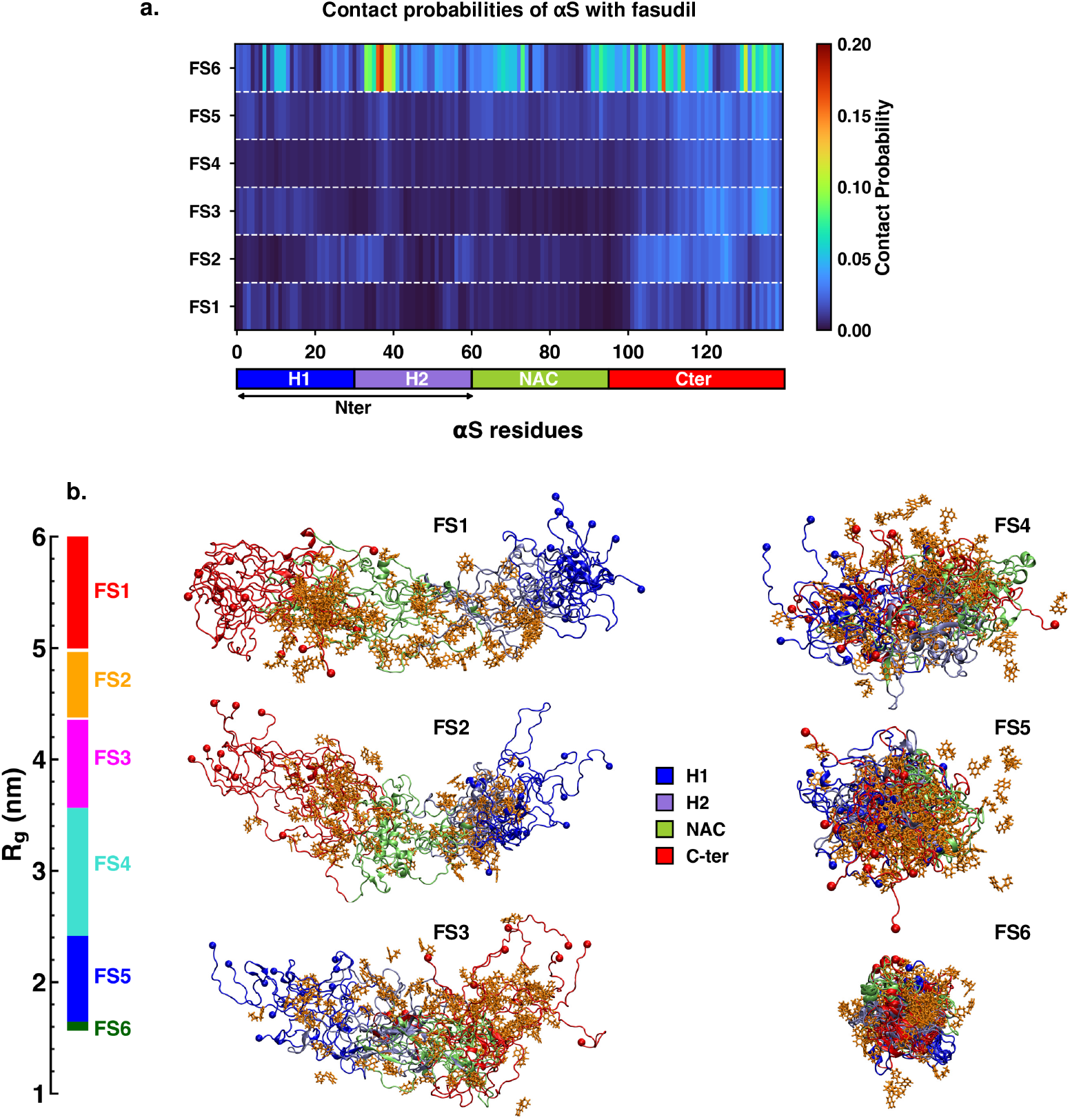
(a) Contact propbability map of residue-wise interaction of fasudil with *α*S. (b) Overlay of representative conformations from the six macrostates (FS1 to FS6) along with the bound fasudil molecules. The conformations are colored segment-wise as shown in the legend. The fasudil molecules are rendered as licorice and colored in orange. The span of the macrostates along the R_*g*_ scale is shown on the right of the snapshots.

The residue-interaction profiles of each of the six individual macrostates of *α*S with fasudil revealed distinct motifs of interaction across the length of the protein, which varies depending upon the relative compaction of the macrostates (Figure 2). In the most compact state FS6, fasudil interacts with the entirety of the protein. The most prominent residue-fasudil interactions are present in the N-terminal and C-terminal regions with charged and polar residues. Fasudil also has moderate interaction propensity with hydrophobic residues (residues 69-75, 92-95) in the NAC region. In the dominant compact state FS5, fasudil interacts predominantly with charged and polar residues mainly tyrosines in the C-terminal region in the range of residues 114, 119 and 121-127, 133 and 136. Interactions are also prevalent in the NAC region as well as N-terminal region (residues 1-14, 28, 35-39) with relatively moderate propensity.

Similar *α*S-fasudil interaction propensities are seen with the C-terminal region in the relatively more extended state FS4. However, the interactions with the NAC regions are suppressed in FS4 compared to FS5 and FS6. With further extension in state FS3, dominant interactions are similarly seen in the C-terminal region and some scattered in the N-terminal residues 1-20. In the extended state FS2, *α*S-fasudil interactions are mainly established with C-terminal residues 104-126 and the N-terminal residues 22-26, 28 and 30-38. Lastly, in the most extended state FS1, the interaction propensities while being lower than other states are restricted to the charged C-terminal region and the H1 (residues 1-30) segment of the N-terminal region. Previous report^29^ has shown that in the bound state, fasudil interactions with *α*S are favorable charge-charge and *π*-stacking interactions that form and break in a mechanism they term as *dynamic shuttling*. The stacking interactions arise from the favourable orientation of fasudil’s isoquinoline ring and the aromatic ring in the side chain of tyrosine residue. Analysis of the time-series of the formation of different intermolecular interactions indicated the formation of charge-charge and *π*-stacking interaction with residues Y125, Y133 and Y136 and the shuttling among these interactions causes fasudil to remain localized to the C-terminal region. Notably, in all the macrostates irrespective of their compaction, fasudil interactions with the C-terminal, especially tyrosine side chains are highly favorable. However, interactions can also be found in the N-terminal and NAC regions. Importantly, as the compaction of the conformation increases, fasudil interactions are increasingly spread across the entirety of the protein, as characterized in these MSM-derived states.

### 4. Structural and kinetic characterisation of metastable states of *α*S monomer in presence of fasudil

Comparison of residue-wise intramonomer contact maps of the equivalent metastable states can be used to identify the differences in the interaction patterns in these states that can be attributed to interactions of fasudil with the monomer. The average inter-residue contact probability maps for each of the six macrostates in the presence of fasudil and three macrostates in water are depicted in Figure 3 and Figure S4, respectively. The residue-wise percentage secondary structure content of the six macrostates FS1 to FS6 are presented in Figure S5. The most exended state FS1 displays weak propensity of long-range interactions between the acidic C-terminal (residues 123-127) and hydrophobic NAC (residues 86-91) regions (see Figure 3a and snapshot i.)). These interactions are formed by *β*-sheet secondary structure among these regions. Interspersed helical structures are also observed in the N-terminal (residues 1-30) and NAC regions (Figure S5). The corresponding extended state MS1 (in water) (see Figure S4 and snapshot i.) shows similar long-range interactions, however spanning more residues interacting via *β*-sheet structures. The newly populated extended state FS2 exhibits similar long-range *β*-sheet forming contacts and helical interactions.

**Figure 3:**
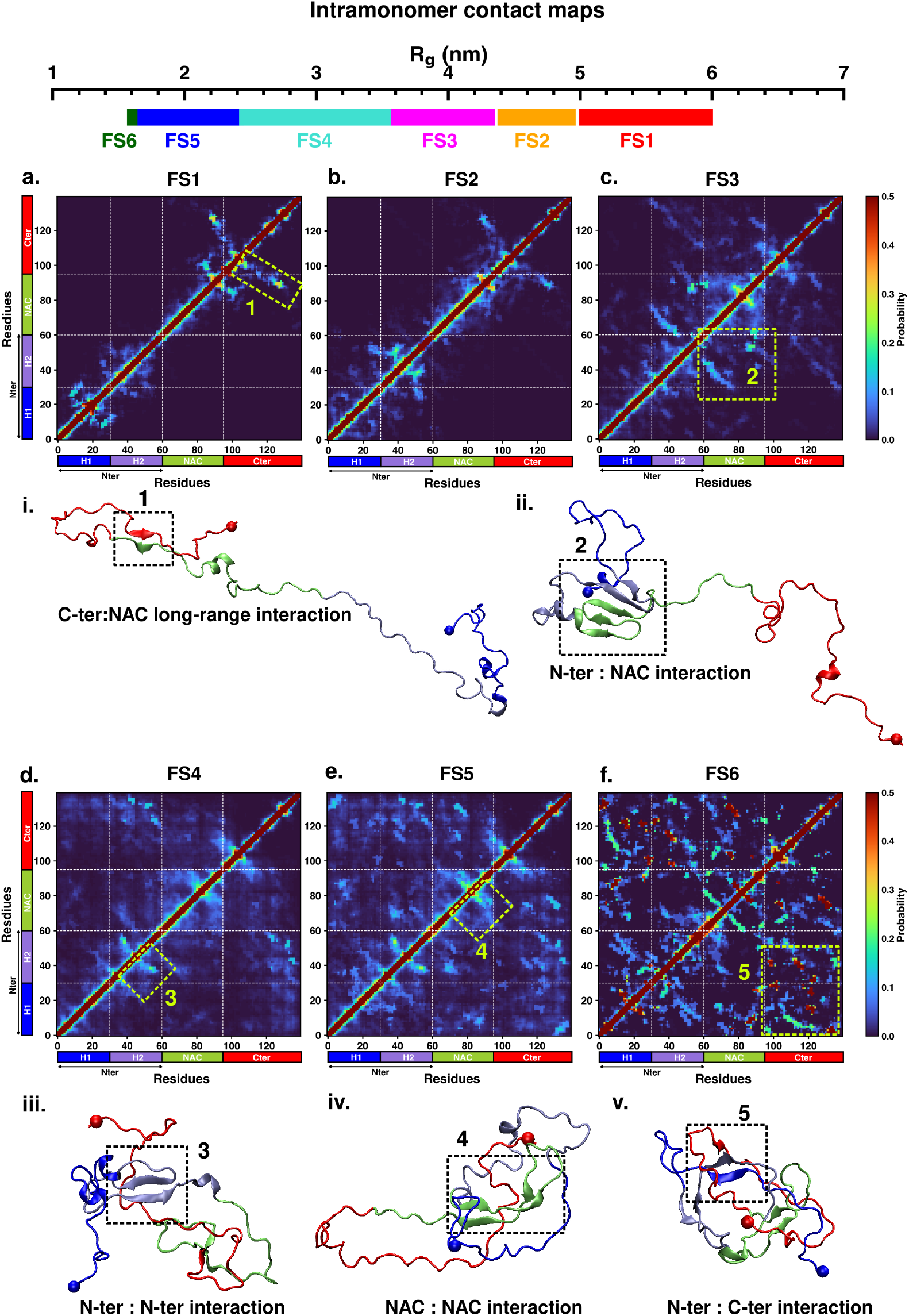
Intrapeptide residue-wise contact probability maps of (a) six macrostates (FS1 to FS6) in the presence of fasudil. Axes denote the residue numbers. The color scale for the contact probability is shown at the extreme right of each panel of maps. The color bar along the axes of the plots represents the segments in the *α*S monomer. The macrostates are marked on the scale of R_*g*_ presented above the contact map panels. Specific contact regions are marked by boxes and numbered. These contacts are illustrated by representative snapshots and the corresponding contacts are similarly marked and numbered.

On the other hand, the new macrostate formed in the presence of fasudil, FS3, that is relatively more compact than FS2, exhibits higher extent of *β*-sheet interactions (*∼*20 %) that is primarily formed in the N-terminal (residues 30-45 and 51-57) and NAC regions (residues 62-80 and 86-95) (Figure 3c and snapshot ii.). In the dominant macrostates FS4 and FS5 that collectively represent the most favourable macrostate MS2 in water, the compact nature is aided by enhanced long-range interactions between the C-terminal and N-terminal (H1 and H2) regions along with contacts formed between the C-terminal and NAC regions as seen in the extended states (Figure 3 d, e). *β*-sheet interactions are also formed among residues within the N-terminal region (see Figure 3, snapshot iii.). On the other hand, in MS2, (Figure S4b) there are relatively fewer long-range interactions and these interactions are mainly between the C-terminal (residues 120 to 130) and the H1 segment (residues 1-30) of the N-terminal region. The absence of contacts between the C-terminal with the H2 segment (residues 30-60) of N-terminal region can be attributed to the *β*-sheet interactions formed between hydrophobic residues in the NAC region and H2 segment. Furthermore, there is a significant propensity of *β*-sheet structures formed within the hydrophobic NAC region in MS2 (Figure S4b and snapshot ii.). This is a key difference between the comparable compact dominant states formed in water and in aqueous fasudil solution.

Finally, the extremely compact state FS6 (more compact than the most compact state in water, MS3) is rich in long-range interactions of the C-terminal with the NAC and N-terminal region forming a network of *β*-sheet interactions. (Figure 3 and snapshot v). Additionally, *β*-sheet interactions are also formed within the N-terminal and NAC regions. The extent of *β*-sheet interactions across the entire length of the protein varies between *∼*20-60 %; the highest propensity is found mainly along residues 37-43 (N-terminal), 75-79 (NAC) and residues 112-116 and 128-131 (C-terminal). Residues extending from the H2 segment of the N-terminal and NAC regions (residues 54-64) form significant extent (*∼*40 %) of helical structures in FS6 (Figure S5f). Overall, with increasing compaction from FS1 to FS6, there is a gradual rise in the interdomain long-range contacts and intradomain interactions manifesting as *β*-sheet interactions. The compaction is further bolstered by helical structures in the amphipathic N-terminal and hydrophobic NAC regions. These distinguishing features among the macrostates formed in water and aqueous fasudil solution are repercussions of *α*S-fasudil interactions that we have explored in the previous section.

The kinetic transition network among the macrostates provides an estimate of the timescales of dynamic conformational transitions (see Figure 4 and Table S1). The timescales indicate that in the fasudil system, the transition from extended state (FS1) to the most populated states of intermediate R_*g*_ (FS5 and FS4) takes place over timescales of 374 and 359 ns, respectively (see Figure 4a-b). In water, however, the equivalent MS1 to MS2 transition timescale is *∼*95 ns (see Figure 4c and Table S1). On the other hand, the reverse transition from the compact metastable states (FS4 or FS5) to most extended states (FS1) occurs over timescales of *∼*35 *µ*s, whereas the comparative MS2 to MS1 transition timescales is nearly *∼*150 *µ*s (see Table S1). In other words, the kinetic conformational transition towards a more compact metastable state is slowed down in the presence of fasudil, while the reverse transition towards extended states is faster. This suggests that small-molecule interaction with *α*S has significant impact on the kinetics of conformational transitions causing a reversal of the general trend of collapse or extension of the macrostates as compared to the dynamics in neat water. To understand the plausible reason for the differential trend observed in the presence of fasudil, we evaluate the transition pathways connecting the macrostates. For the transition from FS1 to FS5 or FS6, 98% of the pathway proceeds via the intermediate states in the order FS2, FS3 and FS4 (Figure 4b). This suggests that fasudil may potentially trap the protein conformations in the intermediate states thus resulting in a longer transition time to attain a more compact state as compared to structural compaction in water. The reverse transition from compact to extended states is faster in aqueous fasudil solution owing to the same reason of passing through multiple intermediate states. The interaction of fasudil with *α*S can result in newer conformational states being more accessible as compared to those attainable in water, therefore hastening the transition when the protein passes through the intermediate partially compact conformations.

**Figure 4:**
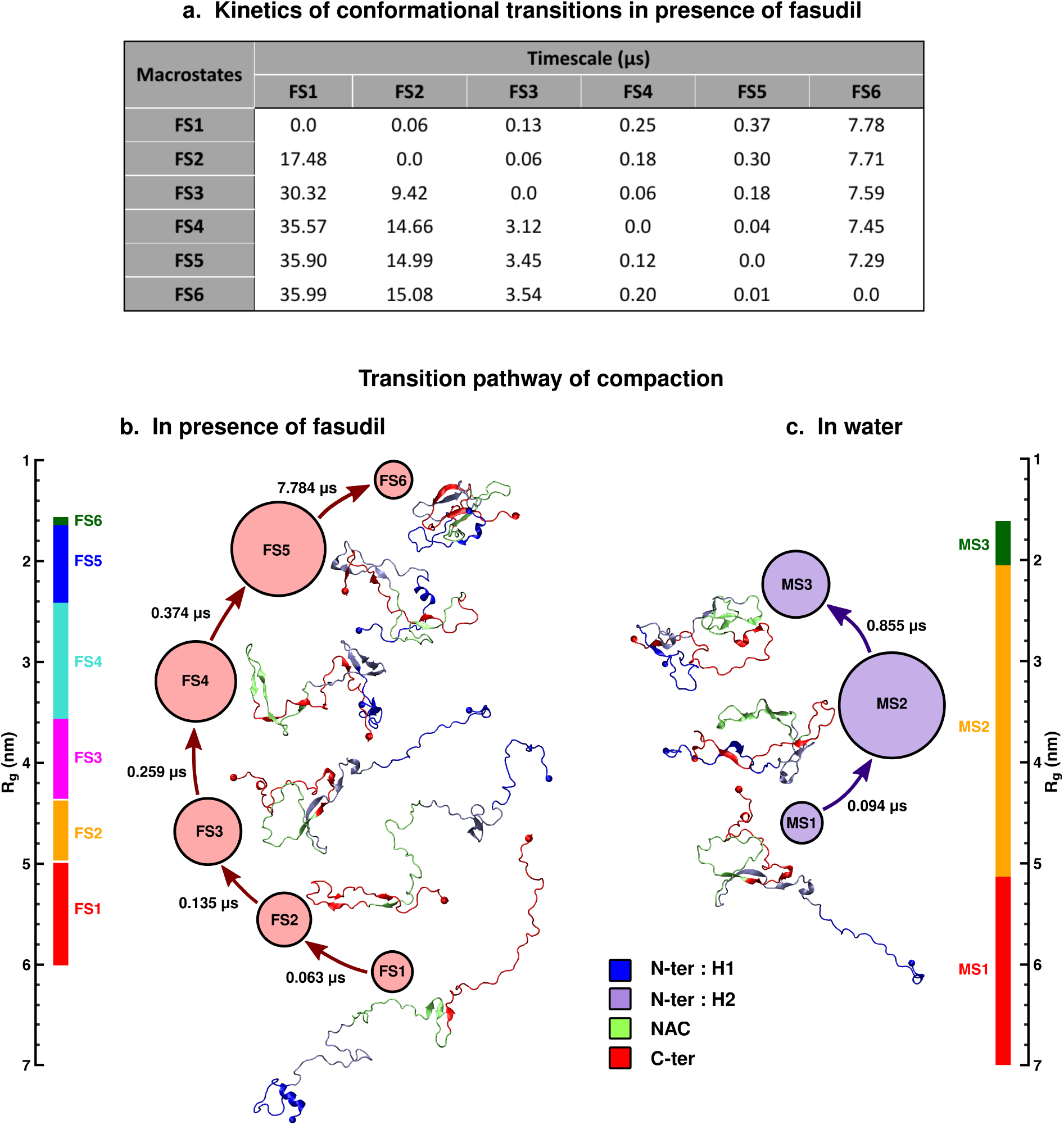
(a) Timescales of dynamic conformational transitions of *α*S in the presence of fasudil. The transition pathway and the associated timescales in the process of compaction in (b) aqueous fasudil solution and (c) water. A representative snapshot for each of the metastable states is presented. The macrostates are marked on the scale of R_*g*_ presented the left and right of fasudil and neat water systems, respectively.

### 5. An entropy expansion mechanism underlies modulation of *α*S conformation by fasudil

The overall analysis of the conformational ensemble of *α*S in presence of fasudil demonstrates that small-molecule binding substantially modulates the ensemble of *α*S. Primarily, there is appearance of newer states of monomeric *α*S in the presence of fasudil relative to that in water. Compared to aqueous conditions, stabilising interactions with the small molecule promotes the formation of both more compact states as well as populates newer intermediate extended states. In general, the conformational ensemble of IDPs consists of distinct conformations that are defined by their occupation probabilities and rapidly interconvert over specific timescales. The behaviour of disordered proteins is thus not determined by a single conformation but is an amalgamation of the conformational properties of the whole ensemble. The binding of small molecules to disordered proteins or regions, therefore, is manifested as modulation of the entire ensemble by shifting the population of the states leading to either an increase or decrease in its conformational entropy.^32,33^ The *entropic collapse* or *folding upon binding* model describes the disorder-to-order transition that IDPs undergo upon interacting with small molecules. This transition leads to the shift in population to a more ordered, predominant state causing a loss of conformational entropy that is compensated by a corresponding enthalpic gain. Conversely, small-molecule binding may also lead to *entropic expansion* in which newer conformational states are populated that are otherwise undetectable.^33^ Additionally, there can be a scenario in which there is no gross change in the conformational entropy and the binding is considered *fuzzy*, ^36^ refered to as *isentropic shift*.

Among the multiple mechanisms of structural ensemble modulation of disordered proteins upon small molecule binding, the observations reported in the present investigation for *α*S monomer plausibly conform with the entropy-expansion model. ^32^ According to this model, the interactions of small molecules with IDPs can promote the exploration of conformations with discernible probability, thereby expanding the disorderedness (conformational entropy) of the ensemble.^20,32,33,37^ Small molecule interaction with *α*S operates transiently, and not strongly as seen in structured proteins. These transient local interactions stabilise the conformations that are inaccessible in the absence of the small molecule. This also has an impact on the kinetic transition timescales leading to either acceleration or deceleration of extension or compaction of states, respectively. This possible mechanism of entropy expansion upon small molecule binding is illustrated schematically in Figure 5.

**Figure 5:**
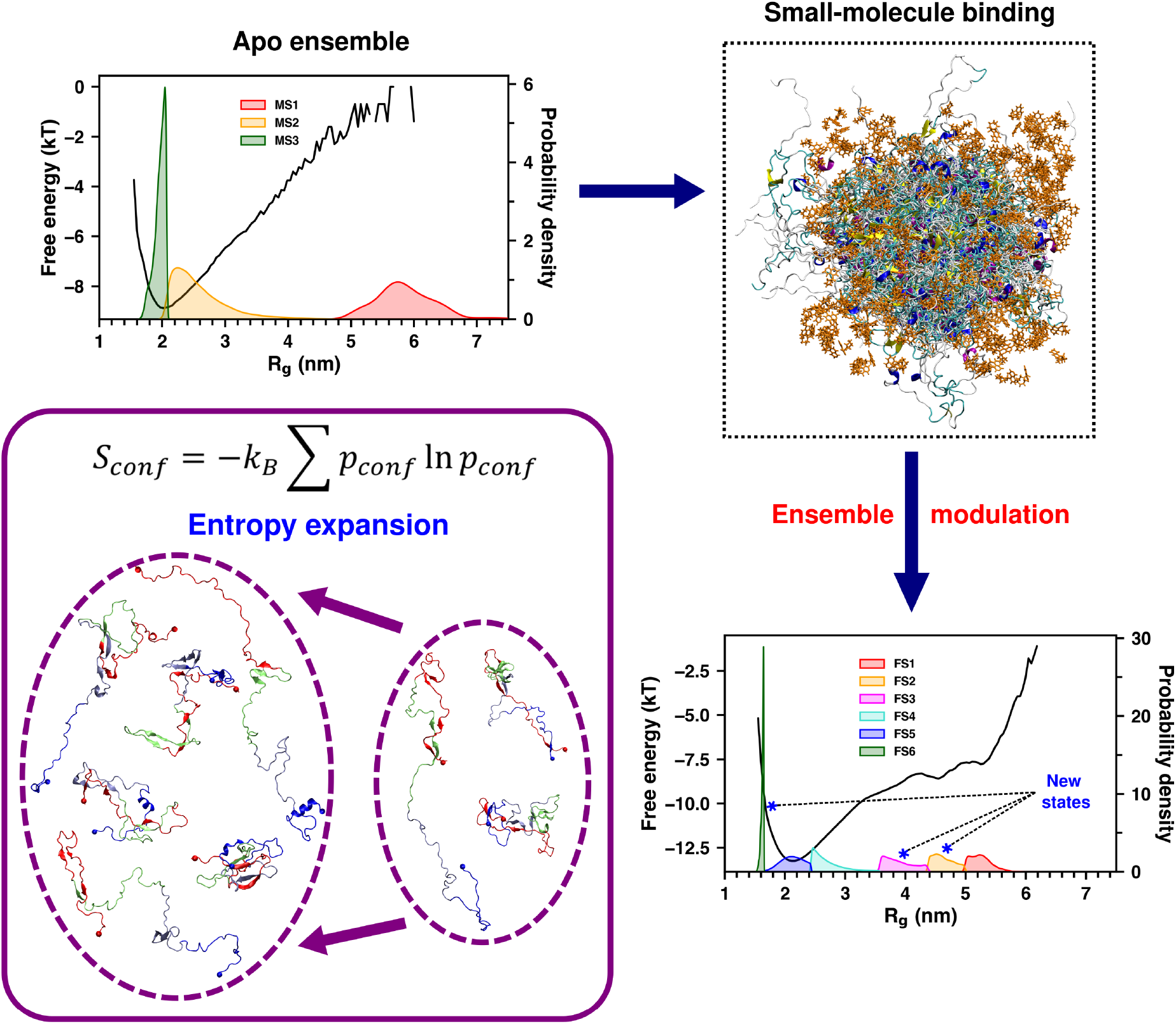
Schematic representation of ensemble modulation by enropy expansion of *α*S monomer by the small molecule, fasudil. The top left panel shows the population distribution and free energy of the three-state ensemble of *α*S in water. The fuzzy ensemble populated in the presence of the small molecule, fasudil is depicted on the right. The ensemble modulation arising from small molecule binding (lower right panel) is shown in the population distribution and free-energy of the increased number of states (6-state model) in the small-molecule ensemble. The new states are marked in the plot. The entropy expansion scheme showing the representative conformations from the metastable states is illustrated in the lower left panel.

### 6. Fasudil hampers *α*S dimerisation process via entropy expansion

Small-molecule (fasudil) modulation of monomeric *α*S ensemble described so far indicated that fasudil interactions with *α*S traps it in multiple conformational states that leads to an overall expansion of the protein conformational space. A previous experimental report shows that fasudil attenuates *α*S aggregation both in vivo and in vitro.^26^ In this view, it would be interesting to investigate the impact that fasudil would have on the early dimerisation process and if the entropy expansion effect seen in *α*S monomer can translate to the self-aggregation of *α*S.

With this motivation, we sought to study the effect of fasudil on early aggregation process of *α*S by simulating two monomers in water and in aqueous fasudil solution (Figure 6a). Three microsecond-long trajectories, in water and 10:1 fasudil:protein were generated and compared. The dimerisation event was monitored by evaluating the minimum distance of approach of the two monomers in water and fasudil solution (i.e. minimum distance between any pairs of atoms between the two monomers). We considered a cutoff of 0.8 nm of minimum distance for the monomers to potentially establish interactions. The probability distributions of minimum distance in the two ensembles are compared in Figure 6b and the time evolution of minimum distance in each of the trajectories in water and fasudil solution are presented in Figure 6c and 6d, respectively. The time profiles indicate that the occurrence of proximity of the two monomers is more frequent in water (see Figure 6c) as compared to that seen in the presence of fasudil in the solution (see Figure 6d). The probability distributions of minimum distance indicate that within 0.8 nm, the intensity of the peak is relatively suppressed in fasudil solution compared to neat water. Evidently, in the simulations, the presence of fasudil and its interaction with the proteins hinders or reduces the propensity of dimerisation as compared to neat water. This observation is consistent with the experimental observation of retardation of *α*S aggregation by fasudil.^26^

**Figure 6:**
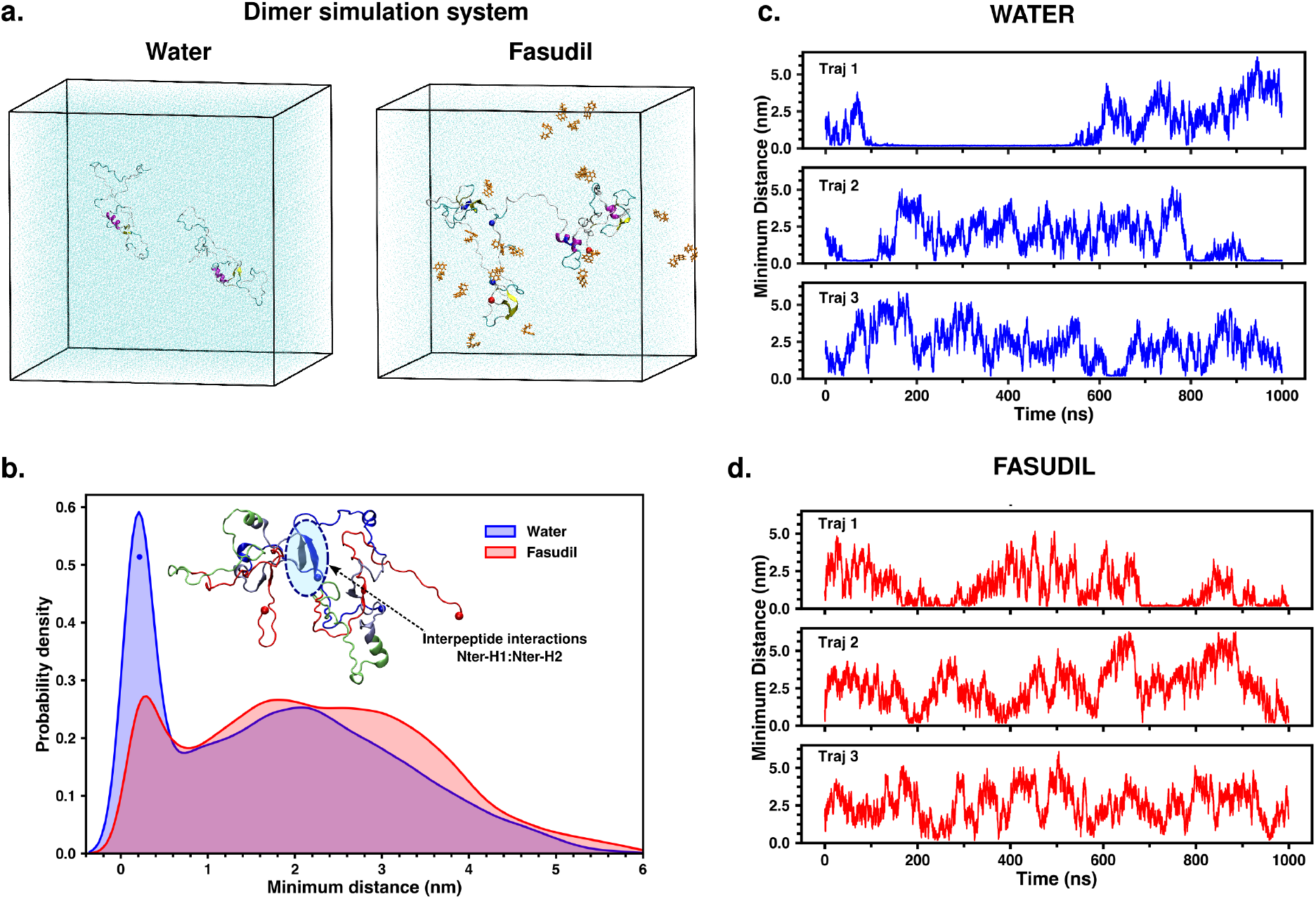
(a) Dimer simulation setup in neat water and aqueous fasudil solution. (b) Probability distribution of minimum distance in the two ensembles. A snapshot of a dimer formed in water is presented and the interpeptide interactions formed between the N-termini of the two proteins is shown. Time evolution of minimum distance of approach of the two monomers in (c) neat water and (d) fasudil solution.

The next logical question concerns the mechanism by which fasudil retards dimerisation and if the process of entropy expansion by creation of new states, as was observed for *α*S monomer, prevails in the process of dimerisation. Towards this end, we first analysed the dimerisation trajectories in water and fasudil solution by clustering the conformations and comparing the clustered states in order to deduce the plausible differences. Both the ensembles, upon clustering (via k-means algorithm, see methods) based on the minimum distance between the two chains, suggested the presence of four metastable states of considerable and competing populations.

Figure 7a and 7d depict the minimum distance distribution of the four clusters in water and fasudil solution, respectively, along with their corresponding populations. The clusters will be referred to as C1, C2, C3 and C4 hereafter, in order of decreasing minimum distance between the chains. State C4 in both the ensembles with minimum distance < 1.0 nm can be considered as the *bound* state in which the monomers are within range of interacting with eachother whereas C1, C2 and C3 are solvent-separated *disaggregated* states. The bound state (C4) accounts for 35.2 % and 25.2 % population in water and in the presence of fasudil, respectively. The interacting sites between the two chains in the bound state C4 can be identified from the inter-monomer residue-wise contact maps as depicted in Figure 7b and 7e. The contact map indicates that in water, the two proteins interact transiently via specific contacts between (a) N-terminal : N-terminal, (b) N-terminal : NAC and (c) N-terminal : C-terminal regions. The snapshots of the bound states in water depicted in Figure 7c show these intermolecular interactions. Some of these dominant interactions are established via *β*-sheets between the N-termini of the two monomers consisting of residues 3 to 8 and 47 to 52. Such *β*-sheet interactions are also formed between the N-terminus (residues 7 to 10) and hydrophobic NAC region (residues 64 to 67). On the other hand, as a signature of reduced dimerisation process in fasudil solution, inter-monomer contacts are highly reduced and present with a small propensity between hydrophobic residues 16 to 19 and 29 to 33 in the N-termini (Figure 7e). However, this proximity does not warrant the formation of secondary structure (*β*-sheet) between these regions as observed in dimers formed in neat water indicating that they do not approach to form stable secondary structures that sustains a dimer conformation.

**Figure 7:**
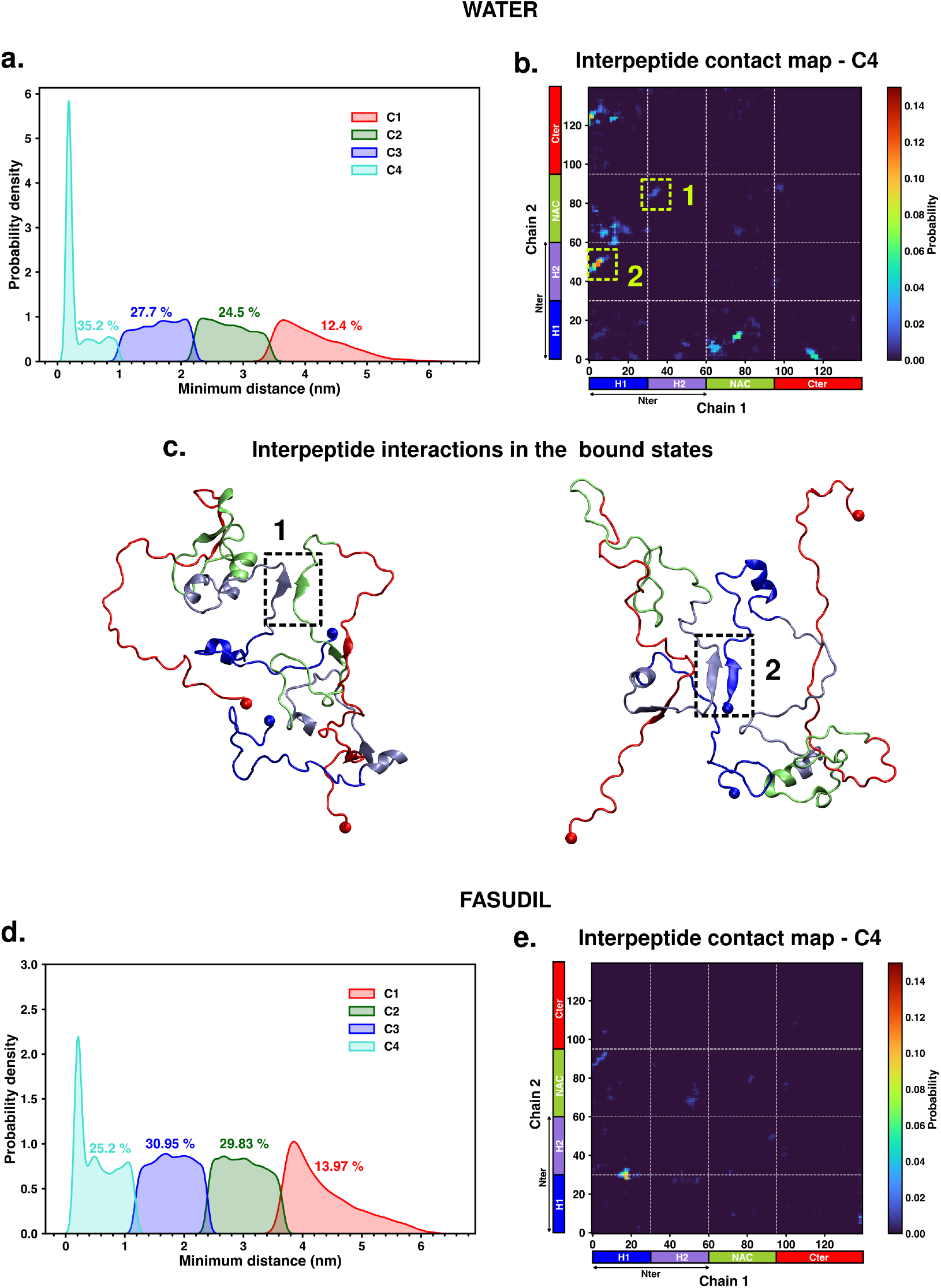
(a, d) Probability distributions of the minimum distance of the four clusters in water and fasudil solution. The percentage populations of each of these clusters is indicated. (b, e) Interpeptide residue-wise contact probability maps of the bound state, cluster C4, formed in water and in presence of fasudil. The axes denote the residue numbers, one axis corresponding to each protein chain. (c) Representative snapshots of the bound state C4 in water and the interpeptide interactions are marked.

We further explored the four conformational states of the ensembles to evaluate if binding of fasudil to the *α*S chains causes entropy expansion in the dimerisation process. For this, we evaluated and compared the extent of compaction of the four states. The probability distributions of R_*g*_ of the monomers in the four clusters in water and fasudil solution ensembles are shown in Figure 8. At the outset, in cluster C4 with monomers in closest proximity, the individual protein chains appear to be more extended compared to the other three states (C1 to C3) in the respective ensembles. However, between the two ensembles, i.e. C4 in water Vs C4 in fasudil solution, the distribution indicates that the protein chains in cluster C4 in the fasudil ensemble are more extended than that in water. On comparing the relative compactness of the disaggregated states (C1 to C3) in the neat water ensemble, the *R*_*g*_ distribution indicates uniform degree of compaction across these three states.(Figure 8a). Notably, in the presence of fasudil, a gradation in R_*g*_ distribution is observed across the states C1 to C3, with the extent of protein compaction progressively decreasing as the monomers move further apart from each other. Together the results from the dimerisation study so far indicate that fasudil apparently impedes *α*S dimerisation and also stimulates systematic diversity in the conformational states of the protein, which essentially alludes to *entropic expansion* as observed when fasudil interacts with monomeric states of *α*S. In the following sections, we further delve into the intra-monomer interactions as well as fasudil binding interactions that govern the observed conformational diversity.

**Figure 8:**
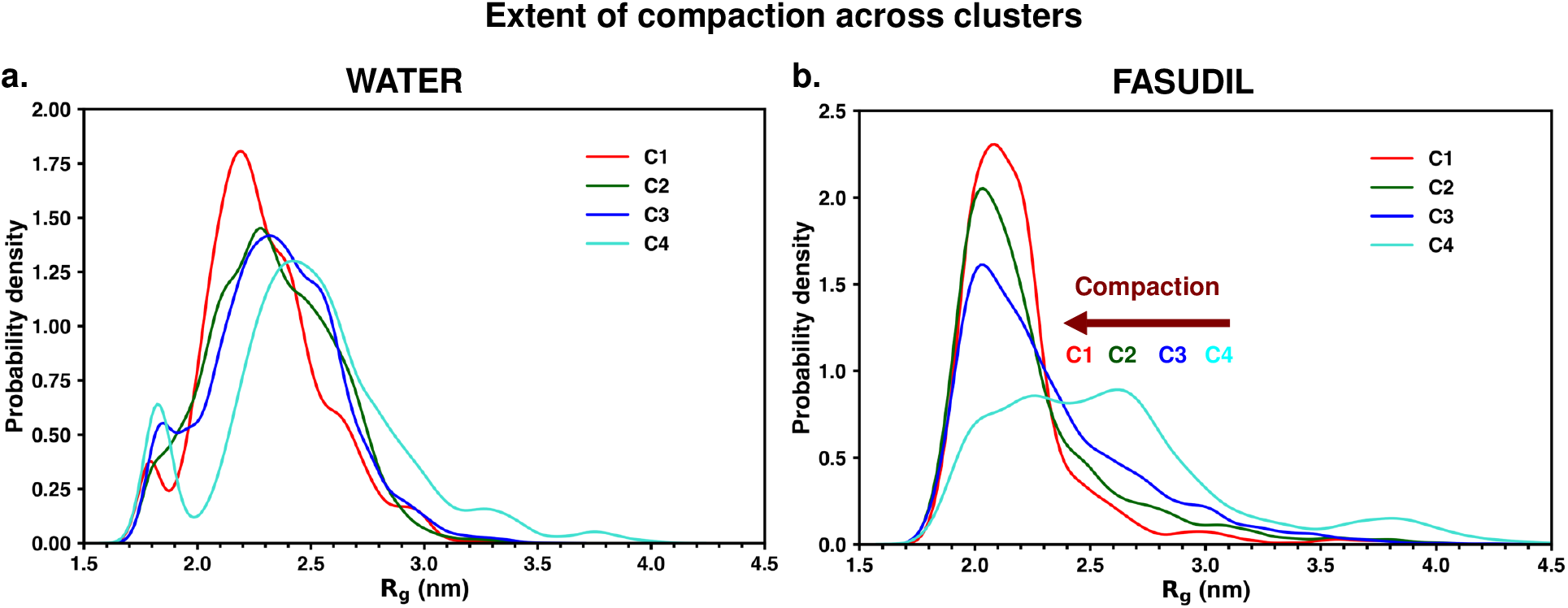
Probability distribution of R_*g*_ of the monomers in the four clusters of dimerisation trajectory in (a) water and (b) in presence of fasudil.

### 7. Preferential interaction with fasudil dictates the extent of dimerisation

The intrapeptide residue-wise contact maps of the states in monomer-dimer equilibrium in water (Figure 9) and fasudil (Figure 10) further elucidate the conformational states and their features that are facilitated by small molecule interactions with the protein. In water, the solvent-separated disaggregated states C1 to C3 display mutually similar intrapeptide contact maps as seen in Figure 9. This substantiates the comparable extent of compaction as observed in these three states (Figure 8a). These states are mainly characterised by long-range interactions of the C-terminus with the N-terminal and NAC region, forming *β*-sheet structures among these regions (see Figure 9 a-c and the adjacent snapshots). The hydrophobic NAC region also forms *β*-sheet contacts with the N-terminal region. However, in the bound state C4, there is a discernible reduction of contacts formed by the N-terminal with the C-terminal region (see Figure 9d). This can be attributed to the observation that in the C4 state, the two protein chains can potentially establish interactions with each other that are mainly mediated by the N-terminal regions of the two chains (see snapshot adjacent to Figure 9d)). This results in the release of long-range N-to-C terminal contacts thus affecting the overall tertiary structure of the individual protein chains leading to extension.

**Figure 9:**
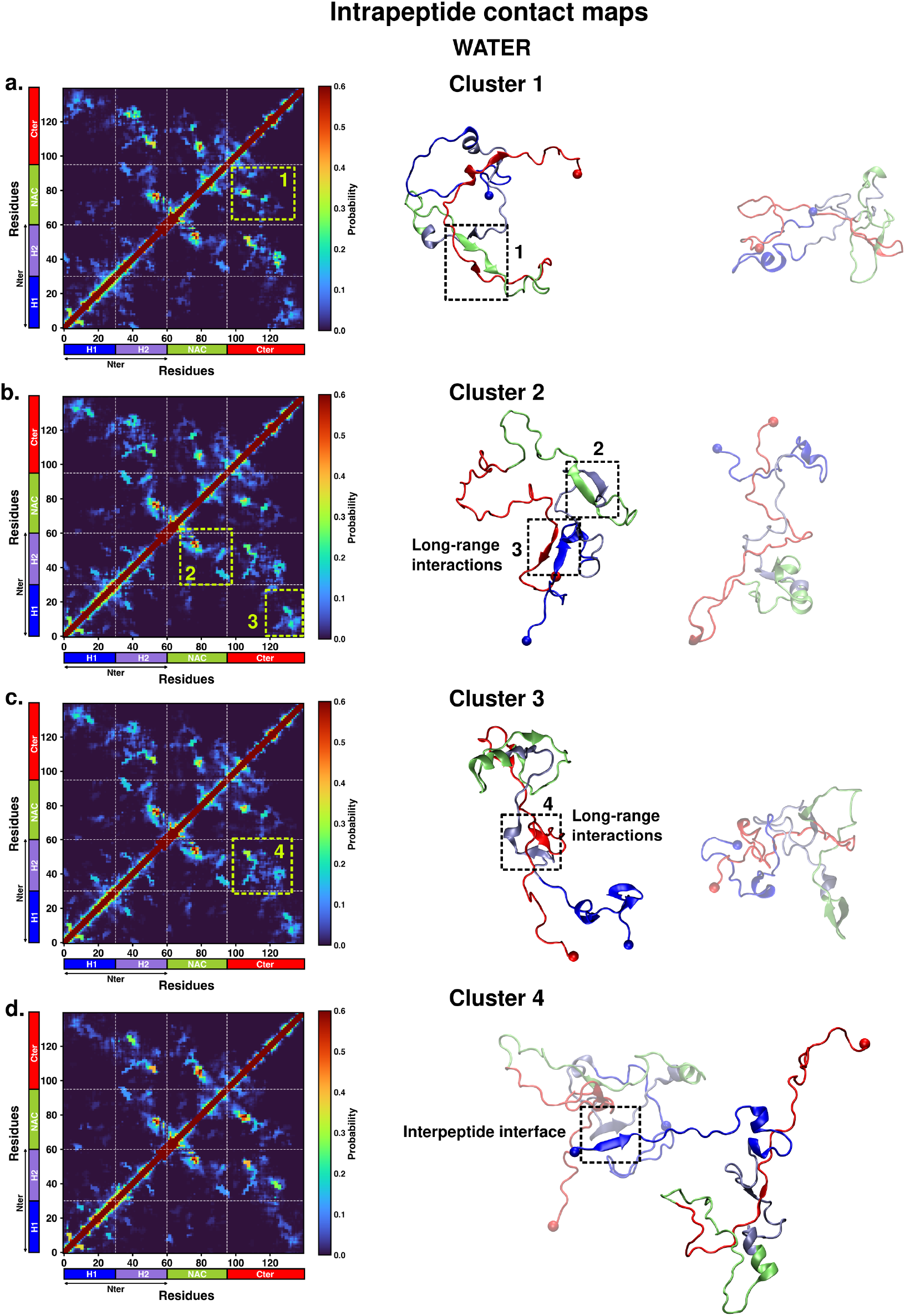
Intrapeptide contact maps of the clusters C1 to C4 (a-d) in water. A representative snapshot of the conformations in each cluster is displayed on the right of the maps. One of the protein chains is rendered opaque while the other chain is shown transparent. Specific contact regions are marked in the contact maps and in the snapshot. The interpeptide interface in the bound state C4 is shown.

**Figure 10:**
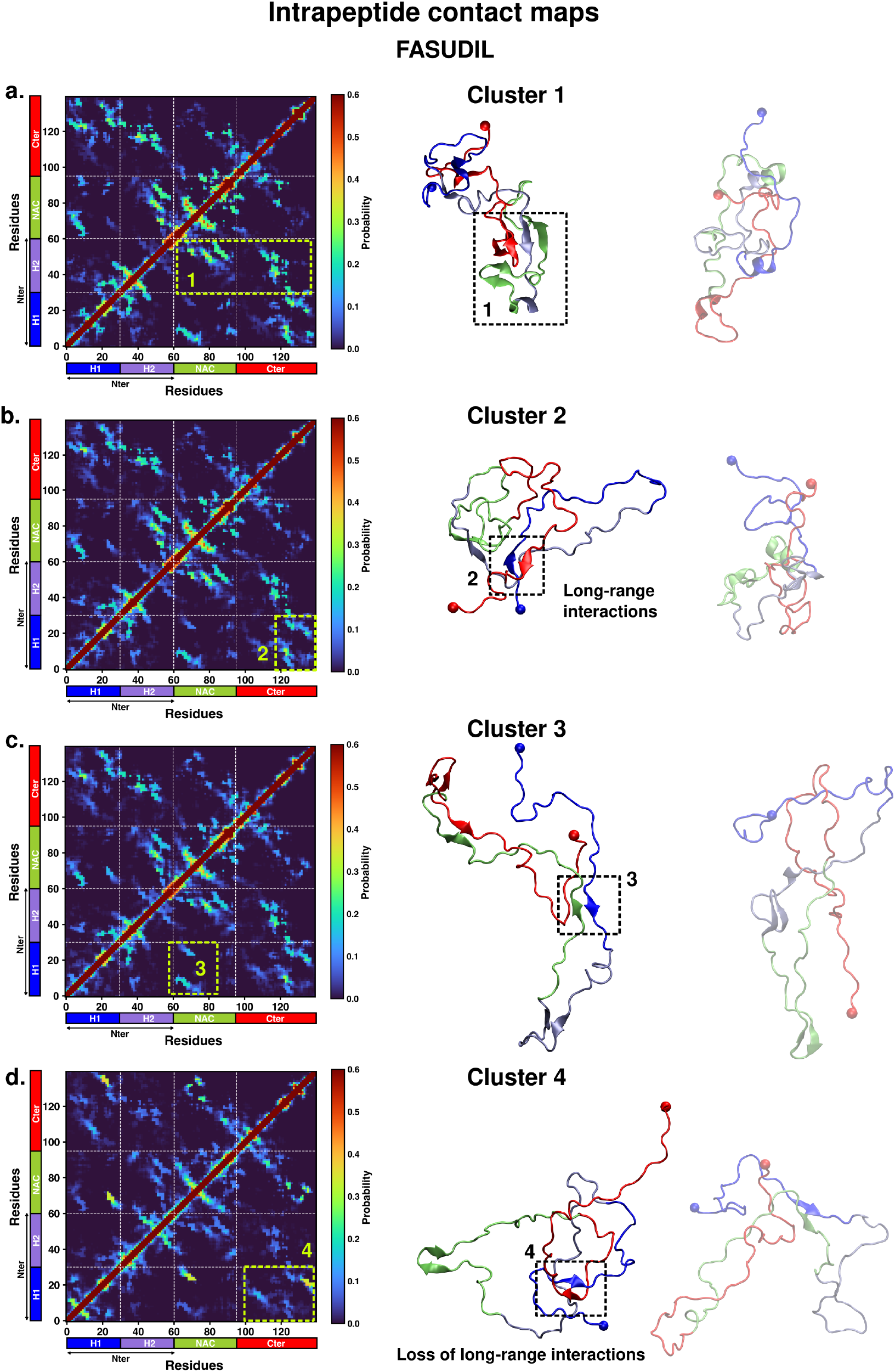
Intrapeptide contact maps of the clusters C1 to C4 (a-d) in the presence of fasudil. A representative snapshot of the conformations in each cluster is displayed on the right of the maps. One of the protein chains is rendered opaque while the other chain is shown transparent. Specific contact regions are marked in the contact maps and in the snapshot.

On the other hand, in the ensemble containing fasudil, the hierarchy in compaction across the states is clearly reflected upon comparison of the intra-peptide contact maps (Figure 10). State C1 (highest minimum distance), being the most compact state (see Figure 8b), has substantial inter-domain interactions including long-range N-to-C terminal interactions, *β*-sheet network within the NAC region and between the NAC and N-terminal regions (Figure 10a and adjacent snapshot). As the states C2 to C4 exhibit incremental expansion, the propensity of the aforementioned interactions gradually reduces (Figure 10 b-d). State C4, deemed as the bound state (protein chains within range of interaction) and being most extended in nature, shows a significant reduction in contacts, especially the long-range contacts between the terminal regions (Figure 10d). While in neat water, the expansion of the chains in the bound state can be ascribed to the loss of long-range interactions between the termini and the concomitant formation of interpeptide interactions; the lack of any stable interpeptide contacts among the protein chains is evident in fasudil solution. Hence it can be postulated that fasudil molecules interacting with the proteins drive the expansion of the chains.

To further ascertain how fasudil promotes conformational diversity in a dimerising system and inhibits dimerisation, we analysed the effect of interaction probability of *α*S residues with all the 20 fasudil molecules present in the solution. A protein : fasudil contact is considered to be formed in a frame when the minimum distance between any heavy atom of fasudil and protein is less than 0.6 nm. The average residue-wise contact probability map of the protein with the fasudil molecules for the states C1 to C4 is depicted in Figure 11. In state C1 (most apart and compact), fasudil interactions with protein residues are prevalent mainly in the charged C-terminal region (Figure 11a). Polar residues such as Y133 and Y136 show significant interaction propensity with fasudil. This is consistent with the experimental report of attenuation of *α*S aggregation by fasudil in which NMR spectroscopy revealed that this effect is mediated by the direct binding interaction of fasudil to specifically two tyrosine side chains, namely Y133 and Y136, in the C-terminal region. ^26^ Furthermore, fasudil significantly interacts with residues 66, 67, 75 to 85, 93 to 95 in the NAC region, comprising of mainly hydrophobic residues interspersed with polar and charged residues. Specific interactions are also formed residues 22, 28 to 39 and 50 to 52 in the N-terminal region. In states C2 and C3 that are more extended than C1, similar protein : fasudil interactions are present but the intensity gradually decreases (Figure 11b-c).

**Figure 11:**
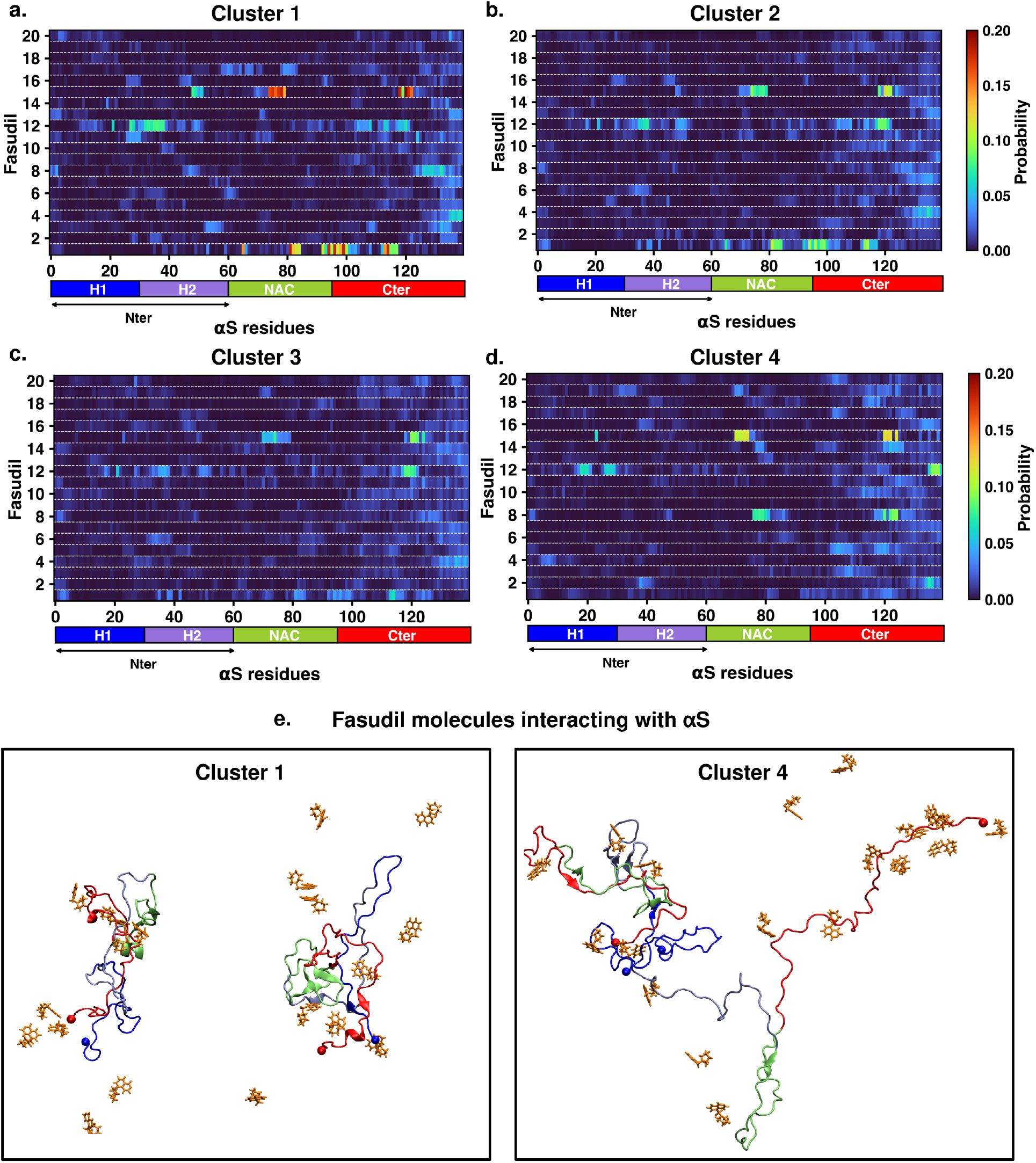
(a-d)Residue-wise contact map of *α*S interactions with the fasudil molecules in the four clusters. The color scale for the contact probability is shown at the right of the map. The color bar along the X-axis of the plots represents the segments in the *α*S monomer. (e) A representative snapshot from C1 and C4 showing fasudil molecules interacting a protein chain.

In state C4, in which the proteins are in closest proximity and most extended in nature, fasudil molecules interact with multiple sites similar to the other states (see Figure 11d) and 11e). In particular, the small molecule strongly interacts with residues 19 to 21, 24, 27 to 30 and 39 in the N-terminal region. In the NAC region, fasudil molecules bind to residues 71 to 75 and 77 to 82. However, interestingly, most residues spanning the C-terminal region interact with fasudil molecules as compared to the C-terminal interactions in other states. It is further noteworthy that multiple fasudil molecules in the solution bind to C-terminal residues simultaneously in state C4 (see 11d) and snapshot in 11e) while fewer fasudil molecules interact with *α*S in the other states C1 to C3. These binding interactions screen the exposure of these residues and thus prevents the formation of interpeptide interactions in state C4 mediated via the terminal regions.

Overall, these analyses indicate that fasudil can interact with specific residues along the entire length of the protein that enables exploration of multiple conformations with varying degrees of compaction. Multiple compact states are favoured in the disaggregated conformation. Also, extended conformations prone to dimerise are also favoured. However, the increased exposure of the residue side chains in these extended states are additionally stabilized by small-molecule interaction that in turn prevents formation of interpeptide interactions and weakens the binding strength of the resulting dimers. Thus, the *α*S dimerisation study in the presence of fasudil substantiates the phenomenon of entropy expansion effect as observed in the monomer and also elucidates the inhibitory effect mediated by small-molecule binding.

## Conclusions

The conformationally dynamic disordered proteins challenge the application of conventional structure-based drug-design methods that relies on the specificity of a binding pocket based on which the drug molecule is designed and optimized. A general framework describes the effect of small-molecule binding to IDPs by modulation of the disordered ensemble by increasing or decreasing its conformational entropy. Here, we characterised the effect of the small molecule, fasudil, on the conformational characteristics of the quintessential IDP, *α*-synuclein. Fasudil has been experimentally reported to curb the aggregation of *α*S both in vivo and in vitro. The ensemble view of long timescale atomic simulations of fasudil binding to *α*S monomer displays a fuzzy ensemble of *α*S with preferential binding of fasudil to the C-terminal region via charge-charge interactions and aromatic stacking. Here, we built a Markov State Model using this data to delineate the metastable conformational states of *α*S and identify the differences that arise from small-molecule interactions. We compared the metastable states of *α*S in the presence and absence of the small molecule fasudil and the model revealed that small-molecule interaction with the protein led to generation of newer states, both more compact and more extended in nature. Moreover, fasudil interacts distinctly with these *α*S states indicating it is the small-molecule interaction that traps *α*S monomer in multiple distinct states, thereby extending the conformational entropy of the system.

We further extended the study to evaluate if small-molecule mediated entropy-expansion also manifests in aggregation by performing *α*S dimerisation simulations in aqueous fasudil solution. The presence of fasudil has a discernable impact by suppressing the dimerisation potential of *α*S. This effect is exerted by creation of multiple conformational states via entropy expansion, as observed in the monomeric system. Structural characterisation of these states indicates that while the two *α*S protein chains are apart from each other, they are maintained in the aggregation-resistant state by maintaining long-range interactions between the terminal regions. Fasudil interacts with residues in the N-terminal, NAC and C-terminal regions in these conformations. Interestingly, it is observed that when the two *α*S chains approach each other, the conformations are relatively more extended owing to the loss of long-range inter-domain contacts between the terminal regions. Fasudil molecules in the solution interact with the exposed charged and polar residues in the C-termini that effectively sequesters these regions and precludes the formation of intermolecular interactions. On the other hand, similar demarcation of conformational states in neat water is not observed, thus reinforcing the observation of entropic expansion arising due to interactions with the small-molecule, fasudil. Furthermore, in neat water, as the proteins approach, the release of the long-range interactions is accompanied by establishing inter-molecular interactions between the two protein chains.

Atomic-level MD simulations have slowly emerged as a a valuable tool for complementing experimental measurements of disordered proteins and providing detailed descriptions of their conformational ensembles.^38–41^ With improvements in force fields^30,42,43^ that model disordered proteins, the accuracy of MD simulations has dramatically increased, as assessed by their agreement with a wide variety of experimental measurements. Integrative approaches use a combination of experimental restraints such as chemical shifts, paramagnetic relaxation enhancement (PRE),^38,39,41,44^ residual dipolar coupling constants (RDCs),^44–46^ small-angle X-ray scattering (SAXS)^47,48^ and computational models from MD simulations to generate a more accurate description of the ensemble of disordered proteins. More recent advances include statistical approaches such as Bayesian inference models like Metainference,^36^ Experimental Inferential Structure Determination (EISD)^49,50^ and Maximum Entropy formulations^51–53^ that also take into account the experimental and back-calculation model errors and uncertainties. Such improved approaches to leverage MD simulations with improved force fields and experimental knowledge-based ensemble generation have shown tremendous potential for describing molecular recognition mechanisms of IDPs with other protein partners; scenarios such as folding-upon-binding^54^ or self-dimerisation. More importantly, a set of recent initiatives have focussed on molecular characterisation of the interaction of small-molecules with IDPs via effective combination of atomistic simulation and biophysical experiments.^20,29^ In this regard, the present investigation takes a step ahead via a quantitative and statistical dissection of the *α*S ensemble in presence of a small-molecule. The implementation of Markov state model in the present work clearly brings out the feasibility of metastable states which would have a finite ‘stationary’ or equilibrium population in presence of small-molecule or in neat water. As a result, the framework provides a direct comparison of entropy of the system in these two environments. Our systematic investigation of the impact of the small-molecule fasudil on *α*S monomer ensemble and the dimerisation process reveals the altering effects of small-molecule interaction on *α*S ensemble by means of entropic expansion mechanism. The retention of this effect inhibiting early dimerisation of *α*S highlights the potential of harnessing small-molecule modulatory effects in drug-design strategies targeting aggregation diseases.

## Methods

### Monomer simulation protocol

A 1500 *µ*s long all-atom MD simulation trajectory of *α*S monomer in aqueous fasudil solution was simulated by D.E.Shaw Research with GPU/Desmond in Anton supercomputer that is specially purposed for running long-time-scale simulations. ^29^ The protein was solvated in a cubic water-box with 108 Å sides and containing 50 mM concentration of Na or Cl ions. Protein, water molecules and ions present in the system were parameterised with the a99SB-*disp* force field.^30^ The small-molecule, fasudil, was parameterised using the generalised Amber force-field (GAFF). Production run of the system was perfomed in the NPT ensemble. Further details of the MD simulation can be obtained by referring to original work by Robustelli et al.^29^ Similarly, a cumulative ensemble of 108 *µ*s (73 *µ*s long continuous trajectory simulation in Anton^30^ and the remaining generated by short simulations in GPU-based servers) of *α*S monomer in water was used for a comparative study with fasudil system.

### Building a Markov State Model of monomer ensemble

In order to identify the metastable states and evaluate the transition kinetics among these states, the two ensembles, namely neat water and aqueous fasudil solution, were used to build a Markov State Model by employing PyEMMA^55^, a popular MSM building and analysis software. First, an appropriate collective variable (CV) or feature describing the ensemble needs to be calculated to build the MSM of a system. Here, the choice of the CV has been done based on the VAMP2 scores (Variational approach for Markov processes)^34,35^ that measures the kinetic variance present in these features and the features can be ranked based on these scores. For this analysis, three features of the system were considered, namely radius of gyration (R_*g*_), inter-residue contact probabilities and backbone torsions. The analysis was performed for both neat water and fasudil system and the scores are presented in Figure S1. The scores indicate that in both the systems, R_*g*_ is the highest scoring metric and therefore most suitable for building the MSM. This is followed by performing a *k* -means clustering of the ensembles on the R_*g*_ feature space into 300 microstates. A transition matrix was then built by counting the number of transitions among the microstates at a specific lag time. To choose the lag time, the implied timescale (ITS) or relaxation timescales of the systems were calculated over a range of lag times and plotted as a function of lag time. The timescale at which the ITS plot levels off is chosen as the lag time to build the final MSM. The ITS plots corresponding to the neat water and aqueous fasudil systems are presented in Figure S2; lag times of 60 ns and 3.6 ns were chosen for neat water and aqueous fasudil systems, respectively. At these lag times, by identifying the gaps between the ITS, a three-state model for neat water and a six-state model for aqueous fasudil system was built. Lastly, the transition path theory^56–58^ was used to ascertain the transition paths, fluxes and timescales in these models.

### Dimer simulation protocol

For dimer simulations, two *α*S chains were placed with center of mass distance greater than 60 Å apart from each other. The chains were solvated in a cubic box that corresponded to a volume of 139.54 Å^3^ after minimization. For aqueous fasudil solution, 20 fasudil molecules are added to the simulation box. The all-atom force field, a99SB-*disp* ^30^ was used to model the protein and solvent used was the a99SB-*disp* water model.^30^ The simulation system details are tabulated in Table S2. A time step of 2 fs and the leap-frog integrator was used for the simulations. The V-rescale thermostat^59^ and Parinello-Rahman barostat^60^ were employed for temperature and pressure coupling (1 bar), respectively. The Verlet cutoff scheme^61^ (1.0 nm cutoff) was used to calculate Lennard-Jones and short-range electrostatic interactions and the Particle-Mesh Ewald method^62^ was used for computing long-range electrostatic interactions. The systems were minimized using the steepest descent algorithm, followed by equilibration in the isothermal-isochoric (NVT) and subsequently in the NPT ensemble with position restraints on the heavy atoms of the protein for 100 ps each. Production runs in the NPT ensemble were then performed for a timescale of 1 *µ*s; three simulations each were performed for the neat water and aqueous fasudil solution systems, respectively.

## Supplemental Information

All supplemental figures described in the article (PDF)

## Acknowledgements

We sincerely thank D. E. Shaw research for providing us with access to long simulation trajectories of monomeric alpha-synuclein in neat water and fasudil solution which seeded the project. We acknowledge support of the Department of Atomic Energy, Government of India, under Project Identification No. RTI 4007. JM acknowledges Core Research grants provided by the Department of Science and Technology (DST) of India (CRG/2019/001219).

