## Supplemental figures for "Small molecule modulates *α*-Synuclein conformation and its oligomerization via Entropy Expansion"

### Supporting Information for Modulation of Conformation and Oligomerization of $\alpha$-Synuclein by Small-molecule Fasudil

Sneha Menon and Jagannath Mondal*

*Tata Institute of Fundamental Research, Center for Interdisciplinary sciences, Hyderabad 500046, India*

,+914020203091

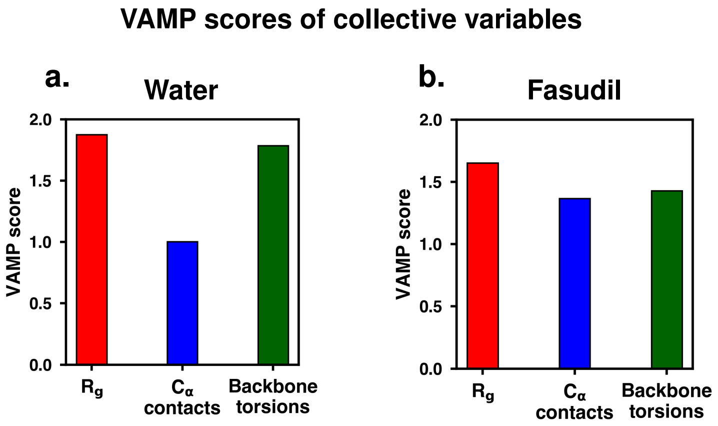


**Figure S1:** VAMP2 score estimated for the features $R_g$, $C_\alpha$ contacts and backbone torsions of the ensemble in (a) water and (b) aqueous fasudil solution.

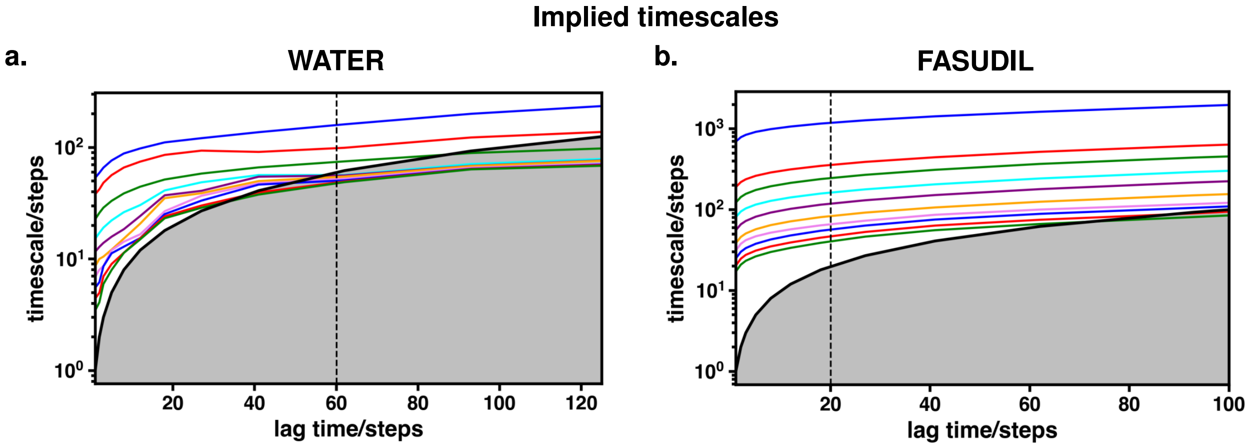


**Figure S2:** Implied time scales (ITS) of the Markov State Model built for the ensembles in (a) neat water and (b) aqueous fasudil solution. The dotted line in each plot marks the chosen MSM lag time for the models.

#### Block analysis of building Markov State Model

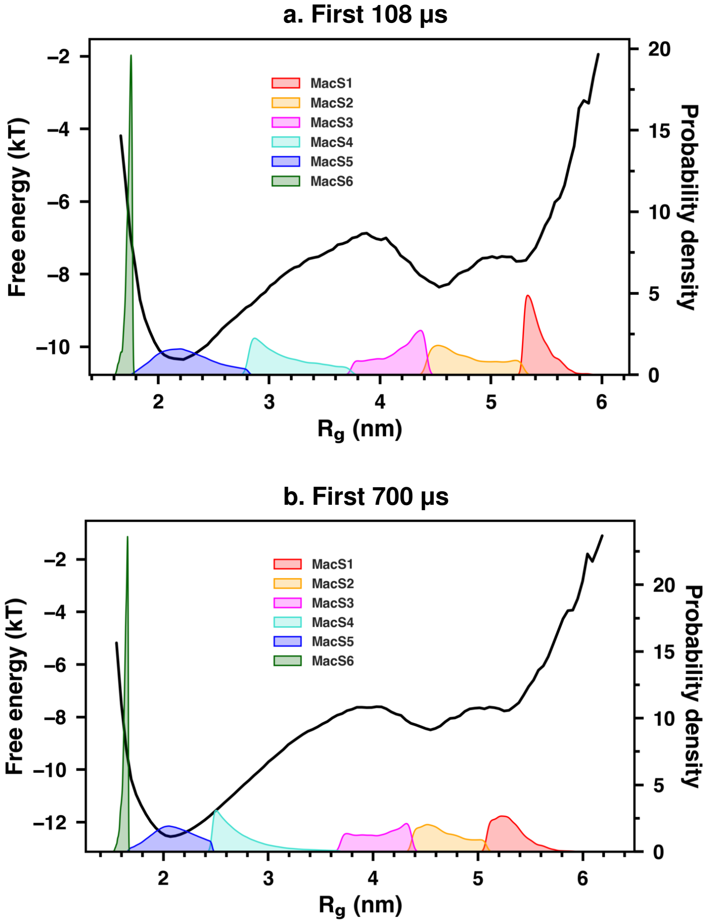


##### c. Macrostate Population

| Macrostate | POPULATION (%) | |
|---|---|---|
| | First 108 μs | First 700 μs |
| FS1 | 1.346 | 0.474 |
| FS2 | 8.926 | 1.295 |
| FS3 | 4.463 | 0.908 |
| FS4 | 9.875 | 16.481 |
| FS5 | 75.051 | 80.320 |
| FS6 | 0.338 | 0.519 |

**Figure S3:** Block analysis of building Markov State Model. Probability density of the macrostates obtained from the MSM corresponding to (a) first 108 $\mu$s and (b) first 700 $\mu$s of the 1.5 ms long trajectory, overlaid with the one-dimensional free-energy landscape estimated as a function of $R_g$. (c) The macrostate populations obtained from the MSM of the two blocks of data.

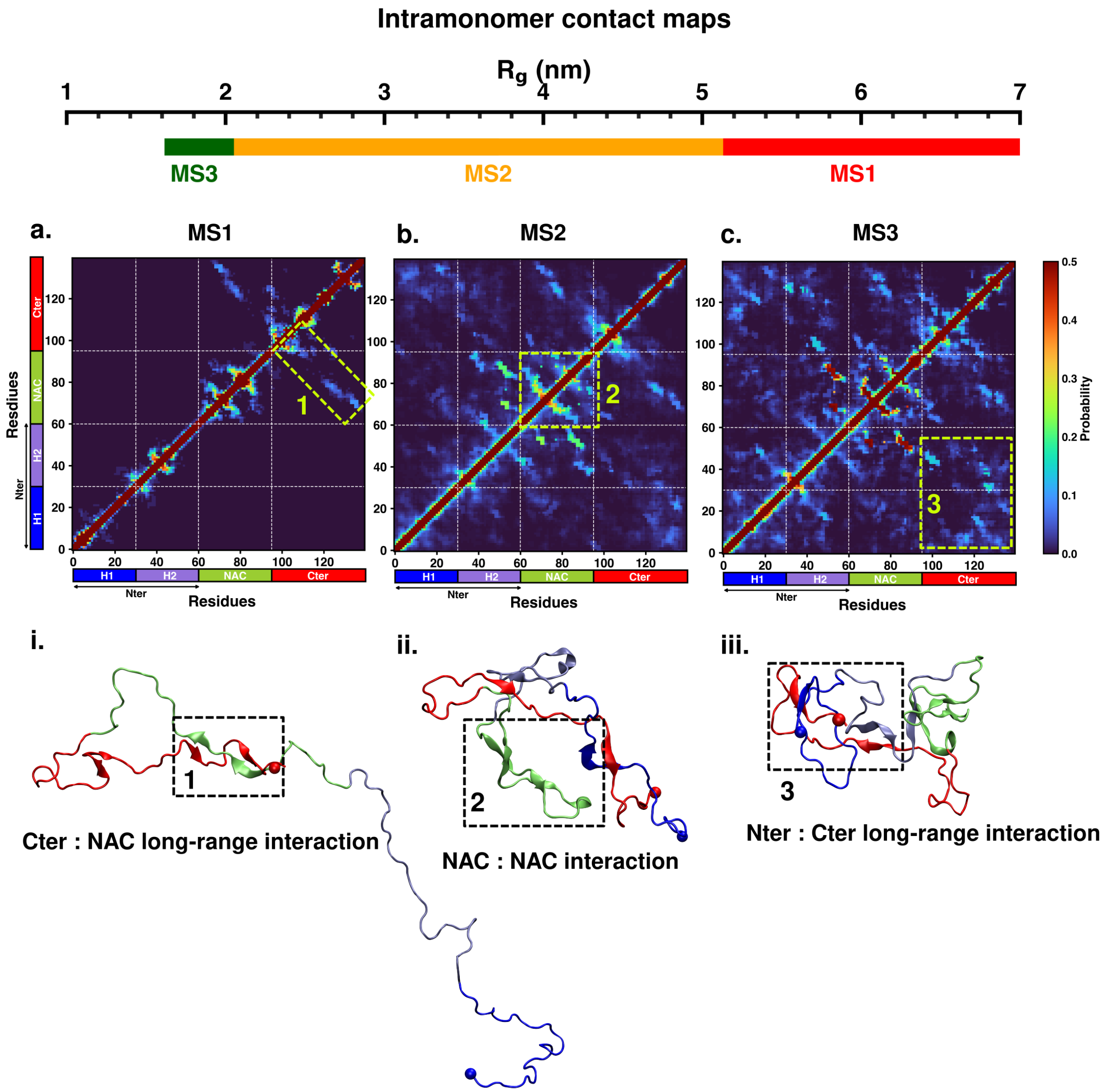


**Figure S4:** Intrapeptide residue-wise contact probability maps of the three macrostates (MS1 to MS3) in neat water. Axes denote the residue numbers. The color scale for the contact probability is shown at the extreme right. The color bar along the axes of the plots represents the segments in the $\alpha$S monomer. The macrostates are marked on the scale of $R_g$ presented above the contact map panels. Specific contact regions are marked by boxes and numbered. These contacts are illustrated by representative snapshots and the corresponding contacts are similarly marked and numbered.

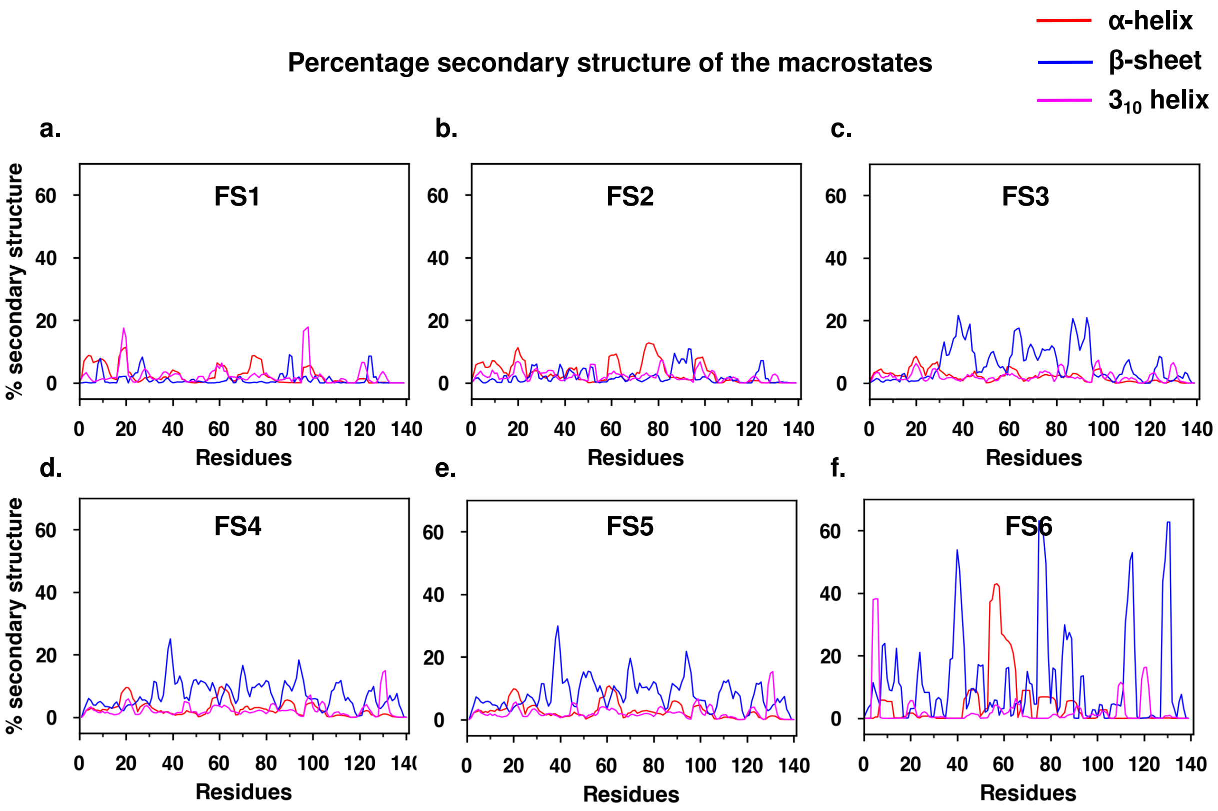


**Figure S5:** Residue-wise percentage secondary structure estimated for each of the six macrostates of $\alpha$S monomer simulated in the presence of fasudil.

**Table S1:** Timescales of conformational transitions of $\alpha$S metastable states in neat water.

#### Kinetics of conformational transitions of α-synuclein in water

| Macrostates | Timescale (μs) | | |
|---|---|---|---|
| | MS1 | MS2 | MS3 |
| MS1 | 0.0 | 0.094 | 1.208 |
| MS2 | 154.9 | 0.0 | 0.855 |
| MS3 | 155.3 | 0.128 | 0.0 |

**Table S2:** Simulation system details of dimers in neat water and aqueous fasudil solution.

| Dimer Simulation details | | | |
|---|---|---|---|
| **System In Neat water** | **Box size (nm)** | **No. of water molecules** | **Total no. of atoms** |
| 3 replicates | 13.954 × 13.954 × 13.954 | 90173 | 357782 |
| **Cumulative simulation time : 3 μs** | | | |

| System In Fasudil solution | Box size (nm) | No. of water molecules | Total no. of atoms |
|---|---|---|---|
| Run 1 | 13.954 × 13.954 × 13.954 | 89992 | 357562 |
| Run 2 | 13.954 × 13.954 × 13.954 | 88179 | 357510 |
| Run 3 | 13.954 × 13.954 × 13.954 | 88155 | 357414 |
| **Cumulative simulation time : 3 μs** | | | |
